# Fast implicit and slow explicit learning of temporal context in cortico-cerebellar loops

**DOI:** 10.1101/2024.08.19.608568

**Authors:** Luca Mangili, Charlotte Wissing, Devika Narain

## Abstract

One is seldom aware of the anticipatory and preemptive feats that the eyeblink systems achieves in daily life but it frequently protects the eye from projectiles gone awry and insects on apparent collision courses. This poor awareness is why predictive eyeblinks are considered a form of implicit learning. In motor neuroscience, implicit learning is considered to be slow and, eyeblink conditioning, in particular, is believed to be a rigid and inflexible cerebellar-dependent behavior. In cognitive neuroscience, however, implicit and automatic processes are thought to be rapidly acquired. Here we show that the eyeblink system is, in fact, capable of remarkable cognitive flexibility and can learn on more rapid timescales than previously expected. In a task where we yoked contextual learning of predictive eyeblinks and manual responses in humans, well-timed eyeblink responses flexibly adjusted to external context on each trial. The temporal precision of the predictive eyeblinks exceeded that of manual response times. Learning of the well-timed eyeblink responses was also more rapid than that for the manual response times. This pattern persevered with the use of a cognitive strategy, which seemed to accelerate both types of learning. These results suggest that behaviors associated with the cerebellar cortex that were previously believed to be inflexible and largely implicit, can demonstrate rapid and precise context-dependent temporal control.

## Introduction

Previous work in cognitive neuroscience has proposed that human decision-making is governed by dual processes: a fast process, which is intuitive, automatic, and relies on associative memory, and a slow process, which is effortful, conscious, and relies on explicit thinking (Stanovich and West, 2000; Evans, 2003; Kahneman, 2011). This appears to reflect daily experiences where we make rapid implicit decisions based on our intuition, whereas, deliberate decision-making often requires awareness and consumes time. Motor function, on the other hand, may require a different approach. Automated motor functioning often requires extensive practice, while awareness could enhance the acquisition of motor skills. Furthermore, both implicit and explicit motor processes may be at play simultaneously (Mazzoni and Krakauer, 2006; Taylor et al., 2014) and may be vulnerable to conscious control to different degrees (Mazzoni and Krakauer, 2006; Benson et al., 2011; Miyamoto et al., 2020). It is, therefore, unsurprising that unlike cognitive decision-making, in motor learning, the acquisition of implicitly learned functions is demonstrably slower than those acquired through explicit learning (Mazzoni and Krakauer, 2006; Huberdeau et al., 2015; McDougle et al., 2015; Miyamoto et al., 2020).

Matters are further complicated when one considers the different neural circuits that have been ascribed to implicit and explicit learning processes. Frontal cerebral cortical circuits are associated with explicit learning (Taylor et al., 2014) and the fast component of motor adaptation (Mazzoni and Krakauer, 2006; Huberdeau et al., 2015; McDougle et al., 2015; Miyamoto et al., 2020). On the other hand, cerebellar circuits are believed to be involved in implicit processes (Taylor et al., 2010; Ito, 2011; Izawa et al., 2012; Taylor and Ivry, 2014) and are often associated with slower learning (Mazzoni and Krakauer, 2006; McDougle et al., 2015; Miyamoto et al., 2020). The view that implicit behaviors could be encoded by the cerebellum is generally consistent with other perspectives in neuroscience (Wolpert et al., 1998; Ito, 2011), and it is generally believed that the implicit component of thinking relies on associative memory processes (Stanovich and West, 2000), which the cerebellum is known to encode (McCormick and Thompson, 1984; Thompson, 1988; Albergaria et al., 2018). Furthermore, recent physiological and anatomical work in rodents has demonstrated that the cerebellum and cerebral cortex are connected in functional anatomical loops (Gao et al., 2018; Chabrol et al., 2019; Guo et al., 2021) and share task-related representations (Wagner et al., 2019). It remains unclear, however, why such disparities in acquisition rates exist across different domains and, given that implicit and explicit processes are ascribed to different interconnected brain regions, can investigations of learning patterns generate testable hypotheses for neural acquisition mechanisms across interareal circuits?

We examined these questions through the lens of temporal perception, specifically interval timing behaviors that contain both explicit and implicit learning readouts, which are attributed to frontal cortical and cerebellar circuits, respectively. We develop a novel task that represents a hybridization of a manual context-dependent timing task, an explicit learning task whose origins were recently traced to the macaque medial frontal cortex (Wang et al., 2018), with classical eyeblink conditioning, an implicit learning behavior that is widely known to be subserved by cerebellar circuits (Medina et al., 2000, 2002; Christian and Thompson, 2003; Heiney et al., 2014; ten Brinke et al., 2015). In general, cerebellar-dependent (McCormick and Thompson, 1984; Ito, 2011) and implicit behaviors(Bond and Taylor, 2015), are often considered rudimentary and inflexible, like many associative learning behaviors that have been studied since the times of Pavlov(Rescorla, 1988). It is unclear whether such forms of conditioning could exhibit context-dependent flexibility in timing, which are behaviors more often associated with computations of the associative cerebral cortex (Wang et al., 2018). On the other hand, cerebellar theories have long extolled the capacity of the cerebellar cortex to enable pattern separation and encode different contexts (Marr, 1969; Albus, 1971; Billings et al., 2014; Cayco-Gajic et al., 2017; Litwin-Kumar et al., 2017; Lanore et al., 2021), although there is no concrete experimental evidence for the realization of this hypothesized function in cerebellar-related behaviors.

Here, we investigate the learning rates of context-dependent temporal responses and measure their relative implicitness and explicitness to determine how these systems are entwined and what insights can be obtained about the underlying neural circuits believed to encode these behaviors.

## Results

### A paradigm to study implicit and explicit learning of temporal context

In a virtual-reality experiment, human participants were shown a contextual cue (Context) in the form of a *Light* or *Dark* tunnel with statistically equivalent but randomized visual configurations on each trial (Figure 1a,b) to avoid the use of spatial information to determine response times. After the Context onset, a brief flash of light (Cue, conditioned stimulus) was presented (Figure 1a,b) to demarcate a starting time at which participants could begin timing. After the onset of the Cue, the tunnel would set into motion at a constant speed for both contexts such that depth information could not be reliably used for the timing. Participants were expected to learn that each Context condition required responses at different times following the Cue through a delayed button press, i.e., the correct manual response time for the *Dark* tunnel was 800 milliseconds after the Cue and for the *Light* tunnel, 1600 ms after the Cue. The Context was randomly sampled on each trial, and if learned correctly, participants were expected to respond flexibly at different times depending on the Context. Learning of these intervals was facilitated using feedback for incorrect responses when the manual response was not made within a time window. Interval timing responses are known to scale with duration, a phenomenon known as scalar variability in timing (Meck, 2003), which is consistent with Weber’s law. We, therefore, used a Weber fraction window around each interval, called an omission window, where responses were permitted as correct. Responses made after the Cue and outside the omission window elicited punitive feedback in the form of a low-latency periocular airpuff, which resulted in a reflexive eyeblink or unconditioned response (UR, Figure 1c left, Supplementary figure 1a,b). The omission window, therefore, refers to the time window where the airpuff was omitted.

**Figure 1:**
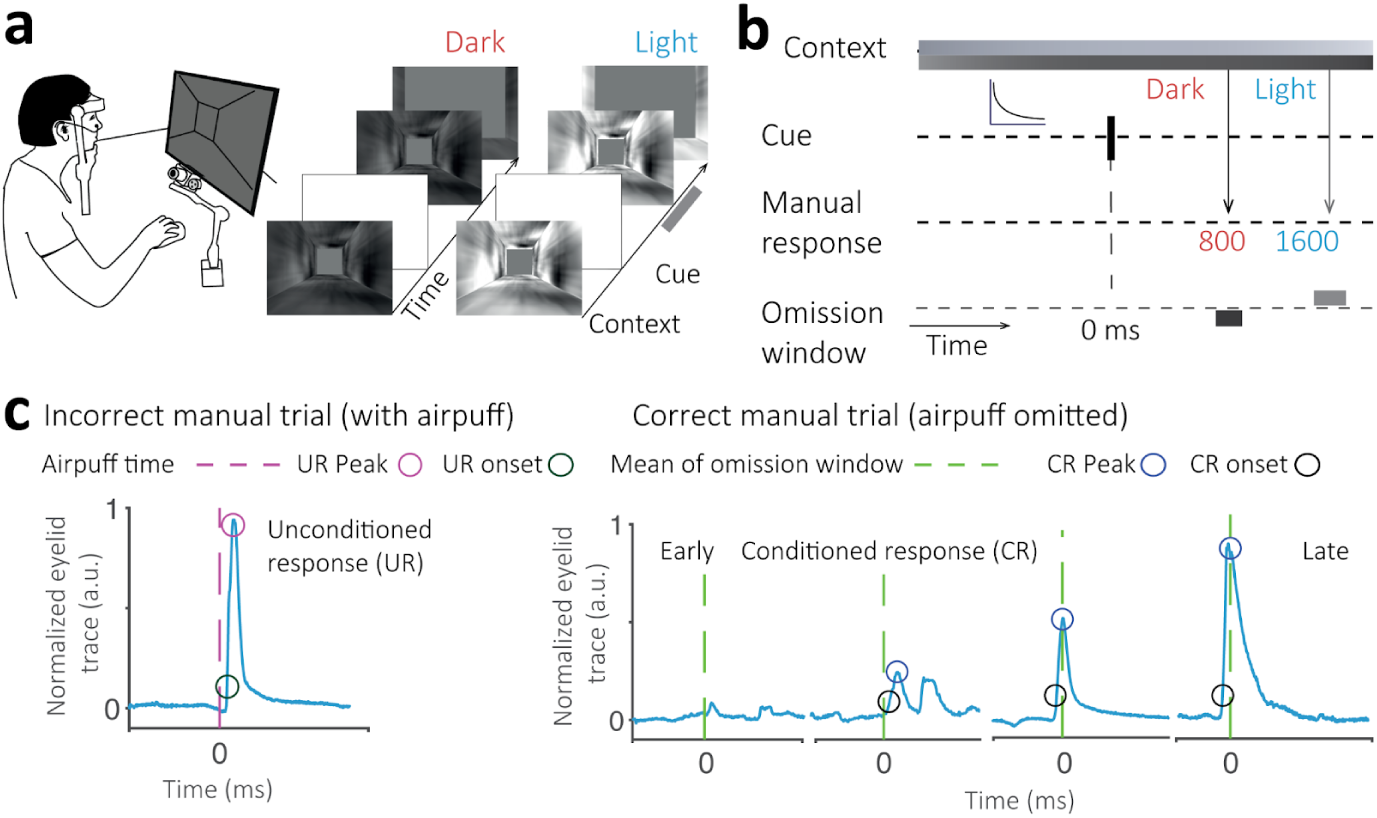
Task design and eyeblink metrics. a-b) Head-fixed participants are presented with a contextual cue, Dark (Red) or Light (Blue) tunnel. Following a transient Cue, participants are expected to generate context-based manual button press responses at 800 and 1600 ms for the Dark and Light context, respectively. On trials where manual timing responses fall outside a Weber-fraction window of the expected times (Omission window), a low-latency periocular airpuff is administered. Eyelid movements are tracked by an infrared camera. c) Left: Administration of an airpuff (purple, dashed line) evokes a reflexive eyeblink or unconditioned response (UR) in the normalized eyelid trace (blue). Right: After learning, we find conditioned responses (CR) or predictive eyeblinks in the absence of the airpuff at the anticipated time of the correct manual response (green dashed line). CR and UR peak (blue/pink) and onset (black/dark green) metrics are also shown.

In this task, the Cue is analogous to a conditioned stimulus (CS), and the paradigm can be considered a form of reverse-trace conditioning. In classical trace conditioning, the airpuff or UR arrives at a single interstimulus interval, and over time, this causes the eyelid to develop a predictive closure response to protect the eye from the airpuff at the anticipated interval(Woodruff-Pak and Disterhoft, 2008). This predictive eyeblink response occurs at the anticipated time even in the absence of an external perturbation like a periocular airpuff. Participants often lack awareness of such a conditioned eyeblink response and cannot control or report its occurrence, thereby making it an ideal implicit measure of learning. To measure awareness, at the conclusion of the main experiments, participants were asked to report awareness for various aspects of the task, including the button press, unconditioned eyeblink elicited from the airpuff and, if present, a conditioned response in the absence of the airpuff.

The manual and eyeblink tasks were coupled such that the airpuff provided punitive feedback following an incorrect manual response (button press at a time outside the omission window elicited a reflexive eyeblink) and the omission window (lack of airpuff and reflexive eyeblink) signaled the desired time interval of response. Learning the manual response required participants to time an interval from the onset of the Cue flash and generate a button press that they were aware of, could control, and could report. We, therefore, used this as an explicit learning measure for the context-dependent time intervals. These implicit and explicit learning measures are linked because the airpuff provided primary feedback for the accuracy of the manual response time. At the conclusion of the experiment, participants reported their awareness of the tunnel context, Cue, button press and whether they blinked on incorrect or correct trials, where the airpuff was present and not present, respectively.

We know that unconditioned stimuli of various kinds, including periocular airpuffs, trigger proprioceptive signals that arrive at both the prefrontal cortical and cerebellar circuitry (Steinmetz et al., 1989; Powell et al., 1996). It is, however, unclear whether a well-timed predictive eyeblink response can be expected to emerge in our task, where the feedback occurs outside a window of the desired manual response time. From a classical conditioning perspective, in this case, there is no predictable manner to protect the eye with a well-timed response.

Surprisingly, in this task, on correct manual response trials, when the airpuff was omitted entirely, we found predictive eyelid responses (conditioned responses, CR) within the omission window (Figure 1c right, Supplementary figure 1a,b). Perhaps even more surprising results were to be found while examining the context-dependent nature of the conditioned response times.

### Context-dependent implicit and explicit learning of time intervals

The timing task presented above has both cognitive and motor aspects and its implicit and explicit readouts are expected to engage the cerebellar and medial frontal cerebral cortical circuits, respectively. Under these conditions, it is unclear whether acquisition timescales for implicit or explicit learning will be faster or slower. Since trace eyeblink conditioning is largely considered an unsophisticated motor timing behavior and cerebellar mechanisms for such behaviors are generally well-understood, conditioned responses may not exhibit precisely timed or context-dependent character and, overall, may be acquired at rates slower than the explicit button press times.

We examined results from a group of participants that received no overt instructions about the context-dependent nature of the experiment before performing this paradigm and were, therefore, called the *naive* group. We found that after learning, naive participants’ manual response times accurately followed Context and their response times were closely associated with those expected for the *Dark* and *Light* context conditions (Figure 2a,b, Supplementary figure 2a,b, N = 30, t(29) = 38.62, p < 0.001). This context-dependent effect in manual response times could also be found for each participant individually (Supplementary figure 2b, Supplementary table 1). The percentage of correct responses (hit rate) significantly improved for most participants over the course of the experiment (Figure 2b right, t(29) = 13.12, p< 0.001) for both conditions (Figure 2b right, *Dark*: t(29) =7.07, p < 0.001, *Light*: t(29) = 12.24, p < 0.001).

**Figure 2:**
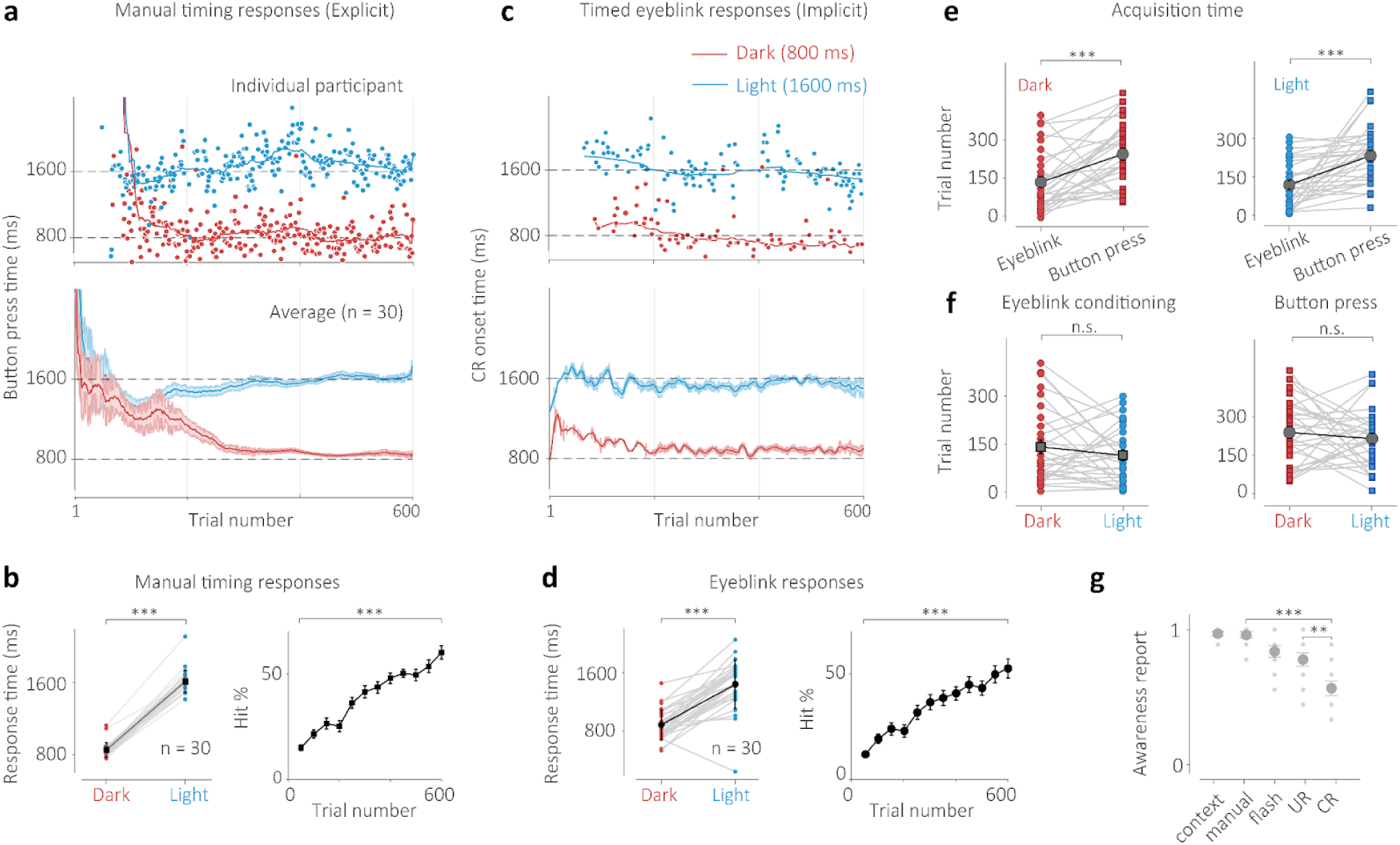
Implicit-explicit context-dependent timing task for the naive group. a) Top: Manual response times of an individual participant for randomly presented Dark (red circles) and Light (blue circles) context trials. Solid lines represent moving averages for each condition. Bottom: Responses averaged across all participants. Error bars indicate the standard error of the mean. b) Left: Average response time for each participant for the Dark (red circles) and Light (blue circles) conditions (trials 301-600). Black squares indicate grand averages and error bars represent standard error. Right: Percentage correct responses (hit rate) progression for all trials (black). Error bars represent standard error. c) Top: Conditioned response (CR) onset time in milliseconds (in the absence of an airpuff on correct manual response trials) for an individual. Color scheme same as a) Bottom: Average CR time across all participants, error bars represent standard error. d) Left: Same as b but for CR onset time and CR percentage (computed as a percentage of all trials in a bin and therefore the measure is correlated with manual hit rate when performance is poor). e) Acquisition time comparison for eyeblink responses (circles) and manual response times (squares) for the Dark (red) and Light (blue) conditions of the same participants. Gray squares indicate grand averages across participants and error bars represent standard error. f) Same color scheme as e) comparing acquisition times within each effector condition (eyeblink and manual response) for the two context conditions. g) Self-reported awareness score (normalized) of participants in a post-experiment survey. Participants rated a 10-point score of how aware they were of the tunnel (context), their manual button press response (manual), the transient flash (flash), blinking after an airpuff on incorrect trials (unconditioned response, UR) or blinking when no airpuff was present on correct trials (conditioned response, CR). Dots indicate individual participants, circles indicate averages, and error bars indicate standard error of the mean.

Remarkably, the conditioned eyelid response onset times (CR time) elicited by participants on correct manual trials also followed context-dependent behavior (Figure 2c,d). Predictive eyelid response times in the absence of an airpuff were indicative of each context’s stipulated time interval and were significantly different for both *Dark* and *Light* contexts (Figure 2d, N = 30, t(29) = 9.77, p < 0.001). This context-dependent timing effect was also found in the CR times of each participant individually (Supplementary figure 3b, Supplementary table 2). The CR percentage, which could only be reliably measured for correct manual trials, also significantly improved over the course of the experiment for most participants and (Figure 2d right, t(29) = 8.39, p< 0.001, Supplementary figure 3c) for both conditions (Figure 2d right, *Dark*: t(29)= 5.68, p < 0.01, *Light*: t(29) = 8.61, p < 0.001, Supplementary figure 3c). This was surprising because the decreased frequency of airpuffs and URs often leads to extinction, however, the manual error rate rarely dropped below 25%, which appeared sufficient to maintain a predictive eyelid response on correct manual response trials.

An important question that arises is whether the predictive eyeblink elicited in the absence of the airpuff was not conditioned in response to the Cue (or CS) itself, in a classical manner, but instead conditioned to the time of the manual button press response on correct trials. If this were true, the conditioned response time would always exceed the manual response time. We analyzed the correlation between the time of the manual response and the onset time of the conditioned response and found no systematic relationship between the two in a vast majority of participants (Supplementary figure 4a,b). Finally, the eyeblink responses often but not always were more accurate than the button press responses (accuracy ratio), however, the precision of these eyeblinks was almost always better than that of the button press responses (precision ratio, Supplementary table 3).

One of the main points of inquiry in this investigation was whether well-timed implicit conditioned eyelid responses would be acquired at rates faster or slower than the acquisition of well-timed explicit manual responses. Using the same algorithmic measures to calculate the convergence of both effector response times, we determined that for the naive participant group, the CR times were acquired significantly faster than the manual response times for the *Dark* (Figure 2e, t(29) = 4.05, p <0.001) and *Light* conditions (Figure 2e, t(29) =4.61, p < 0.01). On the other hand, there appeared to be no significant difference in how rapidly the short (for *Dark*) or long (for *Light*) time intervals were acquired for each effector, viz., eyeblink response (Figure 2f, t(29) = 1.14, p = 0.262) and manual response (Figure 2f, t(29) =0.75, p = 0.459).

Based on earlier work that has investigated the implicit nature of conditioned eyeblink responses(Clark and Squire, 1998; Gerwig et al., 2008), we analysed the self-reported awareness scores (ranging from 0 to 1, 1 is always) for a variety of task-related features (Figure 2g). We found that participants’ awareness of the conditioned eyeblink response was much lower than that for the manual button press response (t(9) = 6.85, p < 0.001). There was also a significantly lower awareness of the conditioned response compared to the unconditioned response (t(9) = 2.92, p = 0.008). These results indicate that the conditioned eyeblink was implicit relative to the awareness of the manual response and the reflexive eyeblink.

### Effect of strategy on implicit and explicit acquisition timescales

To probe how an overt strategy could influence timescales of implicit and explicit learning, we examined a group of participants who were instructed about the context-dependent nature of the timing task and asked to use this information to perform the task accurately (avoid a periocular airpuff); we refer to this group as the *strategy* group. Strategy participants’ manual response times also appeared to accurately distinguish the time intervals associated with the *Dark* and *Light* context conditions (Figure 3a,b, N = 27, t(26) = 32.33, p < 0.001, Supplementary figure 5a,b), which was also the case for each participant individually (Supplementary figure 5b, Supplementary table 4). The percentage of correct responses (hit rate) significantly improved over the course of the experiment for most participants (Figure 3b right, t(26) = 11.45, p< 0.001, Supplementary figure 5c) for both conditions (Figure 3b right, Dark: t(26) = 9.47, p < 0.001, *Light*: t(26) = 9.52, p < 0.001, Supplementary figure 5c), starting at close to 5-10% and stabilizing at over 50%. Here too, the timing of the conditioned eyelid responses elicited by participants on correct trials precisely and often accurately followed the expected context-dependent intervals (Figure 3c,d, N = 17, t(26) = 12.98, p < 0.001, Supplementary figure 6a,b). This was also the case for each participant individually (Supplementary Figure 6b, Supplementary Table 5). The overall CR percentage also significantly improved over the course of the experiment (Figure 3d right, Supplementary figure 6c, t(26) = 9.68, p< 0.001) and for both conditions (Figure 3d right, Dark:t(26) = 8.00, p < 0.001, *Light*: t(26) = 8.23, p < 0.001, Supplementary figure 6c). Moreover, similar to the naive group, the implicit eyeblink responses were acquired earlier than explicit button press time responses for both the *Dark* (Figure 3e, t(26) = 5.11, p < 0.001) and *Light* conditions (Figure 3e, t(26) =4.38, p < 0.001). No significant difference was found in the acquisition times of the two intervals for each Context within the eyeblink conditioning conditions (Figure 3f, t(26) = 0.91, p = 0.37) or the manual timing responses (Figure 3f, t(26) = 1.35, p = 0.19). Similar to the naive group, the precision ratios were less than one, suggesting that the eyeblink responses were more precise than the manual response times (Supplementary table 6). The self-reported awareness scores of the participants in the strategy group indicated that their awareness of the conditioned eyeblink response was much lower than that for the manual button press response (Figure 3g, t(10) = 2.68 p = 0.015). There was also a significantly lower awareness of the conditioned response compared to the unconditioned response (Figure 3g, t(10) = 3.13, p = 0.006).

**Figure 3:**
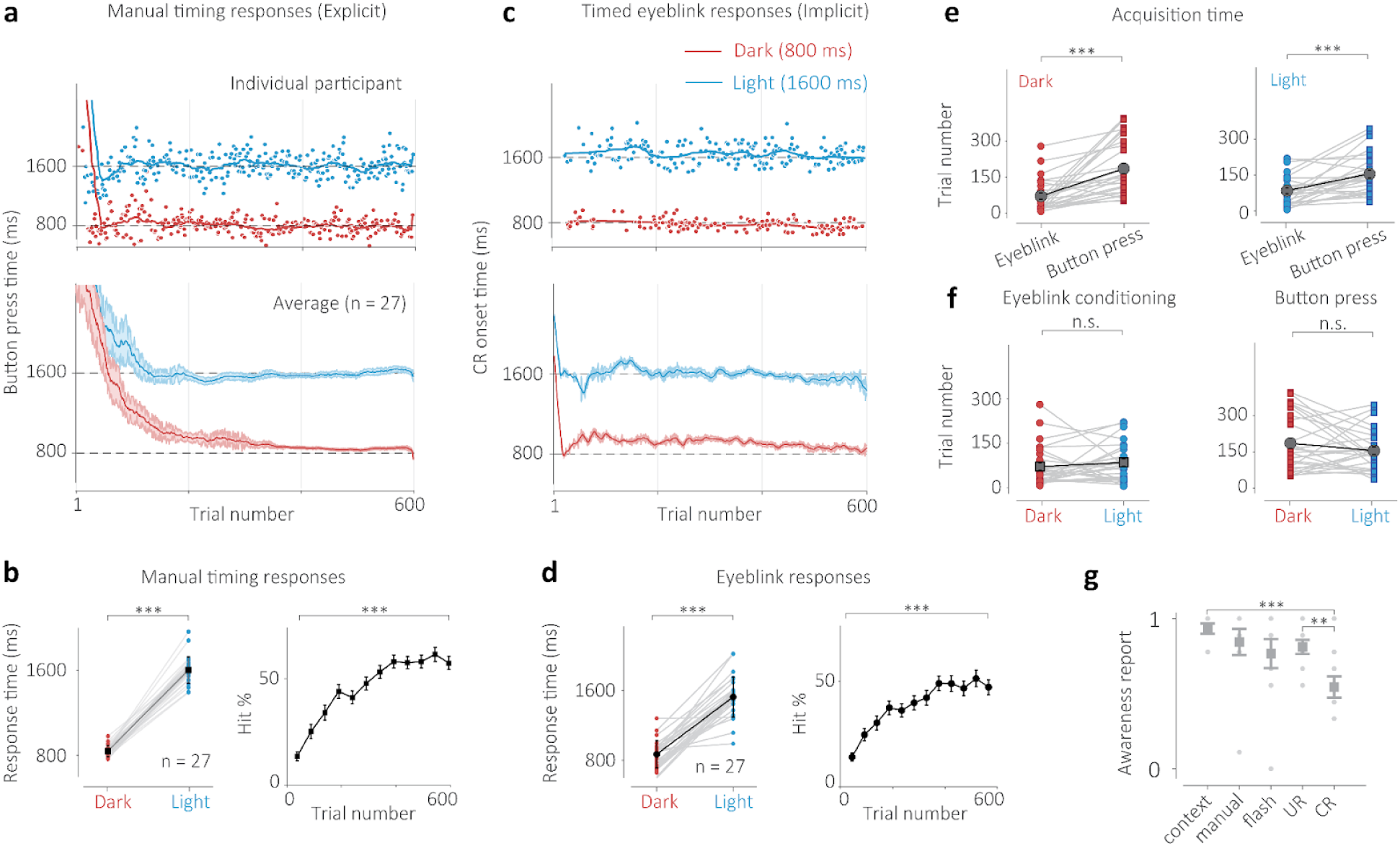
Implicit-explicit context-dependent timing task for the strategy group. a) Top: Manual response times of an individual participant for randomly presented Dark (red circles) and Light (blue circles) context trials. Solid lines represent moving averages for each condition. Bottom: Responses averaged across all participants. Error bars indicate the standard error of the mean. b) Left: Average response time for each participant for the Dark (red circles) and Light (blue circles) conditions (trials 301-600). Black squares indicate grand averages and error bars represent standard error. Right: Percentage correct responses (hit rate) progression for all trials (black) and for each condition (red and blue dots/lines). Error bars represent standard error. c) Top: Conditioned response (CR) onset time in milliseconds for an individual. Color scheme same as a) Bottom: Average CR time across all participants, error bars represent standard error. d) Left: Same as b but for CR onset time and CR percentage. e) Acquisition time comparison for eyeblink responses (circles) and manual response times (squares) for the Dark (red) and Light (blue) conditions of the same participants. Gray squares indicate grand averages across participants and error bars represent standard error. f) Same color scheme as e) comparing acquisition times within each effector condition (eyeblink and manual response) for the two context conditions. g) Self-reported awareness score (normalized) of participants in a post-experiment survey. Participants rated a 10-point score of how aware they were of the tunnel (context), their manual button press response (manual), the transient flash (flash), blinking after an airpuff on incorrect trials (unconditioned response, UR) or blinking when no airpuff was present on correct trials (conditioned response, CR). Dots indicate individual participants, squares indicate averages, and error bars indicate standard error of the mean.

To investigate systematic differences between the naive and strategy groups of participants, we performed a two-way ANOVA to examine the differences in acquisition times for each effector for the two groups for each condition (Figure 4 a,b). We found a significant main effect for the role of strategy for both the Dark (F(1,67) = 9.5, p = 0.003) and Light (F(1,67) = 6.26, p = 0.014) context. We also found a significant main effect for effector response times for the Dark (F(1,67) = 21.89, p < 0.001) and Light (F(1,67) = 34.71, p < 0.001) conditions, which recapitulates the analysis performed showing implicit learning timescales are faster than explicit learning timescales for both groups. Finally, we found no significant interactions (F(1,67) = 0.34, p = 0.56, F(1,67) = 1.04, p = 0.31). Posthoc analysis revealed that using a cognitive strategy reduces the time of acquisition of both explicit and implicit learning measures (Figure 4 a,b).

**Figure 4:**
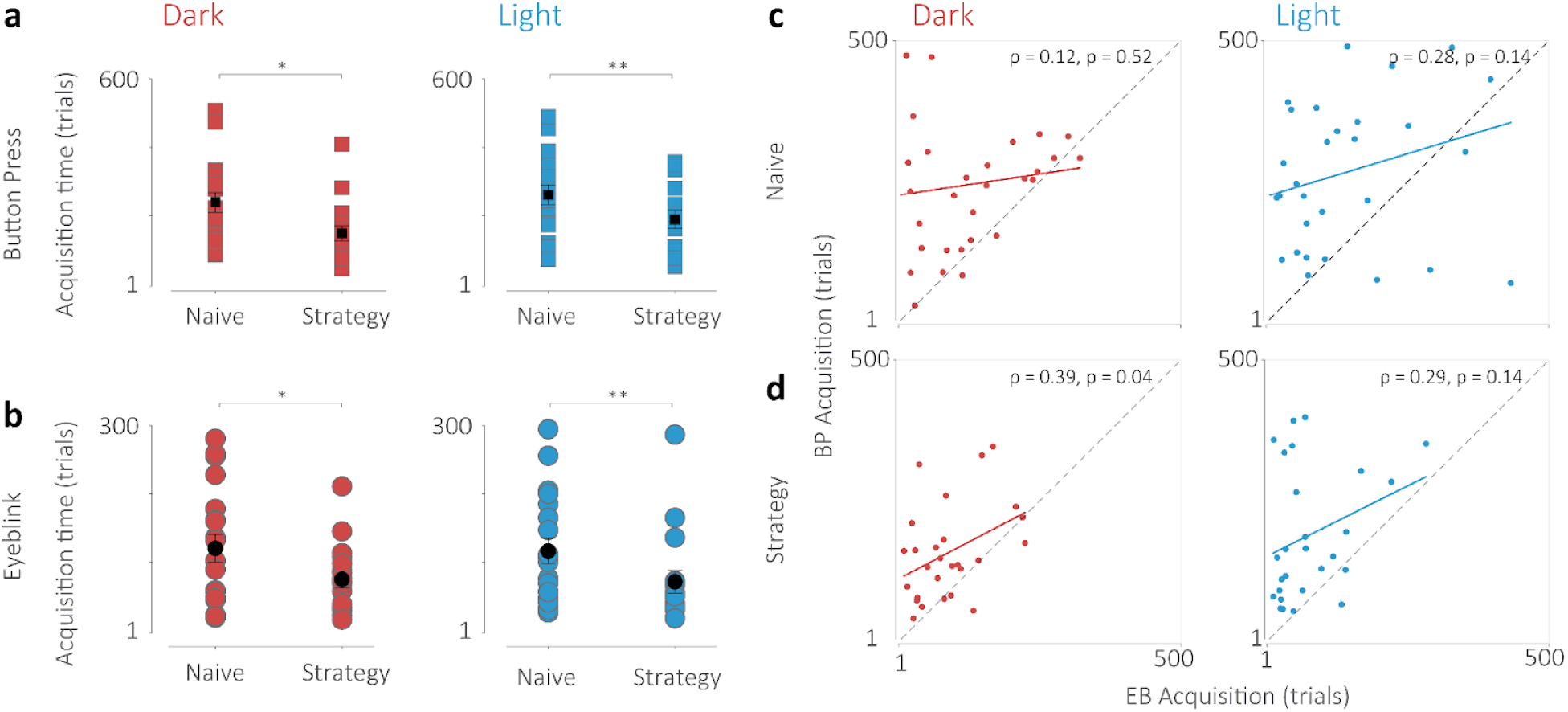
Influence of cognitive strategy on timescales of acquisition of implicit and explicit time estimates. a) Comparison of acquisition times for the manual response times (squares) of the naive and strategy conditions (top) and eyeblink response times (circles, bottom) for the Dark condition (red). b) Same as a) for the Light condition. * indicates p < 0.05, ** indicates p < 0.005. c) Correlation of the acquisition times for the eyeblink and button press responses for the Dark (left, red) and the Light (right, blue) condition for the naive group. d) Same as c) but for the Strategy group. The Pearson’s correlation coefficient and statistical significance are reported for each condition.

To examine whether the eyeblink and manual response learning processes were fully distinct or partly dependent, we plot the correlation of the acquisition times for each participant, condition, and strategy group (Figure 4 c,d). We find that the Pearson’s correlation coefficient does not reveal a significant correlation in any case except the dark tunnel condition for the strategy group (rho = 0.39, p = 0.04). While it is unclear why results are so disparate in this case, we believe these results hint that these processes may not be entirely distinct and may be, at least in part, interdependent.

## Discussion

In summary, we report that the timescales of manual learning of temporal context are relatively slower than those of cerebellar-dependent learning measures of timing. We show that the predictive eyeblink features low on participants’ awareness compared to reflexive eyeblinks and the button press. This suggests that the conditioned eyeblink is relatively implicit compared to the button press in our task but does not completely elude awareness.

Our paradigm was designed to couple explicit learning measures that are attributed to the medial frontal cortex with implicit learning measures long studied in cerebellar circuitry. We also know that cerebellar circuits that are essential for the generation of predictive eyeblinks are disynaptically connected via the thalamus to the motor, premotor, and prefrontal cortical regions in many species. In our task, the only error feedback that enabled learning was the time of the punitive airpuff, which would imply that both the explicit and implicit learning systems would have to utilize this information in order to learn well-timed manual and eyelid responses.

We found that in human participants, both these implicit and explicit measures were able to achieve relatively precise timing for both the *Dark* and *Light* Context conditions, although implicit learning occurred more rapidly than explicit learning of time intervals. This pattern remained unchanged when a cognitive strategy was employed by the participants, although both timescales of implicit and explicit acquisition were accelerated by the use of such a strategy. Previous work in motor neuroscience has shown how cognitive strategies can impact implicit processes antagonistically or synergistically depending on the nature of environmental perturbations(Miyamoto et al., 2020). In these cases, however, the motor effector and, arguably, the motor output circuitry of the implicit and explicit processes may have been shared, thereby enabling resource-dependent juxtapositions of these processes. In our task, the readouts ensue from two different effectors and learning measures are believed to be acquired in two disparate but interconnected neural systems. Nevertheless, results from our correlation analysis show that while these processes are mostly independent, some dependencies may exist. Under these circumstances, one might expect the cognitive strategy to disproportionately advantage the explicit learning system. We found, however, that strategy had a synergistic effect on the acquisition of implicit learning as well. This raises the question of how implicit behaviors like eyeblink conditioning might be vulnerable to overt strategy.

Eyeblink conditioning is frequently performed with a single interval(Christian and Thompson, 2003; Koekkoek et al., 2003; Heiney et al., 2014; ten Brinke et al., 2015), although the acquisition of multiple intervals has shown to be possible(Mauk and Ruiz, 1992; De Zeeuw et al., 2023). Furthermore, voluntary blinking in humans has been studied and elucidates new possibilities for this seemingly rudimentary effector system, which can also use volition to precisely control its output(Rasmussen and Jirenhed, 2017). We believe that our results show a different facet of this behavior, where awareness is low, but the eyeblink exhibits sensitivity to random changes in the context.

Given the speed of acquisition of a well-timed conditioned response based on visual context, we hypothesized that contextual information arrives at the cerebellum directly from primary visual sensory streams, which project to the pontine nuclei. Previous work in classical trace conditioning, however, hypothesizes a role for the prefrontal cortex in funneling extracerebellar afferents that provide a sustained input that bridges the gap between the conditioned and unconditioned stimuli to enable learning of time intervals through long-term plasticity mechanisms(Woodruff-Pak and Disterhoft, 2008; Kalmbach et al., 2009; Siegel and Mauk, 2013; Siegel et al., 2015). Although it has also been argued that despite differences in the input stages, both delay and trace conditioning have common cerebellar mechanisms(Halverson et al., 2018). Furthermore, the hippocampus has also been implicated in classical trace conditioning(Weiss et al., 1999), although patient H.M., in whom the hippocampus was radically excised, was able to generate conditioned eyeblinks for subsecond intervals for both trace and delay eyeblink conditioning(Woodruff-Pak and Steinmetz, 2007). The involvement of other brain regions in this task, therefore, remains unclear and more work is needed to elucidate the role of extracerebellar inputs in the generation of rapid and flexible temporal associations.(Christian and Thompson, 2003; Koekkoek et al., 2003; Heiney et al., 2014; ten Brinke et al., 2015).

Another important consideration is that here we study the subsecond-second range (up to 1600 ms), whereas, cerebellar circuits are known to often support second to sub-second computations(Christian and Thompson, 2003; Breska and Ivry, 2016). Although longer timescales may also be cerebellar-dependent in timing tasks(Ohmae et al., 2017), whether the cerebellum leans heavily on other areas to carry out such temporal computations remains an open question.

## Methods

### Participants

A total of 57 participants (31 female, mean age 23.74 years) provided informed consent to participate in the context-dependent timing task that we developed, in which the error feedback was provided by a periocular airpuff and contextual cues indicated the desired time interval to be generated on each trial. The naive group comprised 30 participants (16 female, mean age 23.77 years), where no explicit information was provided about the context-dependent nature of the correct manual button press responses. Participants were instead encouraged to learn by trial and error based on the airpuff feedback. The strategy group comprised 27 participants (15 female, mean age 23.76), where participants were informed about the context-dependent nature of expected manual button press responses and asked to use this information to generate their responses. Eight participants were excluded from consideration because of one or more of the following reasons: 1) they did not learn the correct button press time for one or both conditions (Hit rate below 20%), 2) they exhibited unconventional unconditioned eyelid responses in the presence of the airpuff (were unable to reflexively close their eyes in response to the periocular airpuff), 3) the spontaneous blink rate of participants was high and interfered with the evaluation of eyelid responses during trials. Participants had no known neurological conditions and gave written informed consent prior to participation. All protocols were consistent with ethical approvals obtained from METC (Medische Ethische Toetsings Commissie) at Erasmus University Medical Center, Rotterdam. No statistical methods were used to predetermine sample sizes, but the sample sizes are similar to those reported in similar studies in other domains.

### Setup

During the experiment, participants’ head positions were fixated using a chinrest. Participants wore a head-mounted custom-made frame that housed a periocular airpuff delivery system. The air pressure system (MPPI-3 pressure injector, Applied Scientific Instrumentation, USA) was housed in a separate room to avoid auditory cues and was activated by a 5V pulse from a low-latency pulse generator (PulsePal2, Sanworks, USA), which triggered pulses after inputs from the CPU of the main computer with microsecond latency. Visual stimuli were displayed using a 240 Hz 23-inch LCD monitor (ASUS VG279QM). A high-speed infrared camera (Basler ace acA640-750um, Basler, Germany) and an infrared lighting system (IR Illuminator TV6700) were trained on the right eye of the participants. Eyelid responses were calibrated to register eyelid movements at the start of the experiments and monitored throughout to ensure the maintenance of the eyelid trace. Participants responded by pressing a low-latency button connected to the CPU by a USB 3.0 port. The experiment was designed using the BonVision package of the Bonsai framework(Lopes et al., 2015) to deliver stimuli and enable low-latency closed-loop control of the input/output streams.

### Stimuli and protocol

The visual stimulus constituting the tunnel contextual cue was constructed from a combination of 8-bit grayscale images (modified from ref(Saleem et al., 2013)). The Dark and Light tunnel images utilized a gamma correction of 1.4 with an overall mean pixel intensity of 53.8 (SD 1.12) and 176 (SD 3.17) for the Dark and Light tunnel stimuli, respectively. These resulted in the stimuli having a Michelson contrast ranging from 0.9-1.0. After the onset of the tunnel context, a random time elapsed (sampled from an exponential hazard function with a time constant of 1 second and truncated at 3 seconds) before a transient flash was introduced to indicate the start of the expected time interval. For this, using a rapid fade-in-fade-out effect, a uniform white image briefly covered the screen for 100 ms. We refer to this flash as the conditioned stimulus (CS) and to the presence of the tunnel context, as a context cue.

The experiment consisted of three sessions with a total of 600 trials where the Dark and Light contexts were presented with uniform probability. At the start of each trial, a stationary tunnel, either Dark or Light, was presented. After a random interval, a brief flash (CS) appeared, which indicated the start of the timed interval and corresponded with the translation of the tunnel in a forward direction. Tunnel wall images for each condition were randomized during placement to eliminate spatial cues that could aid participants in timing while maintaining the statistical properties of the images. Participants made a response using a button press. If the response time lay within a 10% Weber fraction window of the context-specified time interval (800 ms for Dark and 1600 ms for Light), no periocular airpuff was administered otherwise a low-latency airpuff was administered resulting in reflexive closure of the right eye, known as an unconditioned response (UR). Eyelid closures were measured by an infrared camera, calibrated, and registered using the BONSAI(Lopes et al., 2015) framework. After the end of a trial, a new trial started after a random interval drawn from a uniform distribution ranging from 1000-3000 ms.

#### Naive and strategy groups

Participants in the Naive group, at the onset of the experiment, were given no explicit instructions or strategies and remained naive towards the context-dependent nature of expected time intervals. Participants in the strategy groups were given information about the context-dependent nature of the expected time intervals of manual response and were asked to use this knowledge to their benefit in the experiment. A subset of participants in both groups (Naive = 11 and Strategy = 10) were asked to fill out a questionnaire at the end of the experiment.

#### Questionnaire post-experiment to test awareness

At the conclusion of the experiment, participants who were asked to fill out a questionnaire responded to ten questions overall. Five questions required a subjective response asking them to report their awareness of task features on a scale of 1-10, where 1 indicated never noticing the property and 10 indicated always noticing it. These properties, in random order, pertained to how often they noticed whether the tunnel was dark of light (context), whether they were aware of pressing the button (manual), whether they noticed a flash on the screen during trials (cue flash), whether they noticed that they blinked after an airpuff (UR on incorrect trials), and whether they noticed that they blinked when no airpuff was present (CR on correct trials). They subsequently answered five true and false questions to assess their awareness of stimulus contingencies. These asked whether the tunnel was circular in shape (false), whether the patterns of the tunnel walls were the same (false), whether the darkness or lightness of the tunnel had an impact on the correct time (true), whether light and dark tunnels were presented at random across trials (true), whether the white flash was only presented on a few trials (false). The performance on the true and false questions, which was generally high, was used to screen the general awareness of participants to determine their eligibility for the awareness score. The subjective scores were added and normalized to generate an average awareness score (0-1) across participants.

### Analysis

#### Manual response task

The button press times were recorded by BONSAI(Lopes et al., 2015) for each trial relative to the CS onset and were further analyzed offline using a custom script in MATLAB R2022a (MathWorks, Natick, USA). We analyzed the button press times for each individual (using a smooth average to track learning within-condition, Figure 2a, 3a), the percentage of correct trials for each condition within 50 trial bins, and the average response times during the second half of the experiment (trial 301-600) were evaluated as box plots. Responses were considered outliers if they exceeded three times the standard deviation of all responses. The learning traces of button press times for each condition were averaged across subjects to obtain estimates of learning progression within each group. We used the derivative of button press times to evaluate when learning asymptotically converged for each participant. When the derivative was close to zero and learning had reached the Weber fraction window of the expected response for each condition, the trials at which this was achieved were taken as the acquisition time for that condition and participant.

#### Conditioned response analysis

On incorrect manual response trial, administration of the airpuff resulted in the reflexive closure or an unconditioned response (UR) of the eyelid, and after learning, on correct trials, when the airpuff was omitted, a predictive or conditioned response (CR) of the eyelid could be found on a large number of trials. We measured the baseline distribution of eyelid closure prior to the conditioned stimulus (flash) and classified CRs as responses that exceeded three times the standard deviation of the baseline in the epoch following the CS. Note that CRs were only evaluated on trials where the airpuff was omitted (correct manual response trials). The full blink range was computed as the difference between the baseline and full closure measured from the peak unconditioned response time. Drift and fatigue of the eyelids were corrected over a moving window throughout the course of the experiment. The onset time of the CR was computed using the derivative of the normalized eyelid traces when they crossed a participant-specific threshold. CR peak was taken as the maximum eyelid closure within a contiguous predictive eyelid trace (Figure 1c). CR onset times were corrected for systematic biases and were plotted over the course of the experiment to evaluate learning using moving averages. These were then averaged across participants to evaluate the development of CR onset time for the two groups (Figure 2c, 3c). The percentage of CRs was evaluated for each condition within 50 trial bins. Since CRs are only evaluated on correct manual press trials, CR percentage is correlated with hit rates but can diverge at higher hit rates. The average response times were evaluated as box plots over the full course of the experiment. Responses were considered outliers if they exceeded three times the standard deviation of all responses. We used the derivative of CR times to evaluate when learning asymptotically converged for each participant and each condition within the Weber fraction window.

### Statistical tests

The data distributions were assumed to be normal in all cases. Paired t-tests (two-tailed in all cases) were used to evaluate within-group button press (Figure 2b, 3b), CR times (Figure 2c, 3c), hit rates, and CR percentages (Figure 2d, 3d), acquisition times (Figure 2e,f 3e,f) and the model context-dependent response (Figure 5f). The same was applicable to the within-subject analysis of context-dependent manual and eyeblink responses (Supplementary figure 2,3,5 and 6), where corrections were made for multiple comparisons. A two-way ANOVA was used to evaluate the effect of strategy on button-press or CR onset time acquisition for the Dark and Light context conditions (Figure 4).

---

**Supplementary figure 1:**
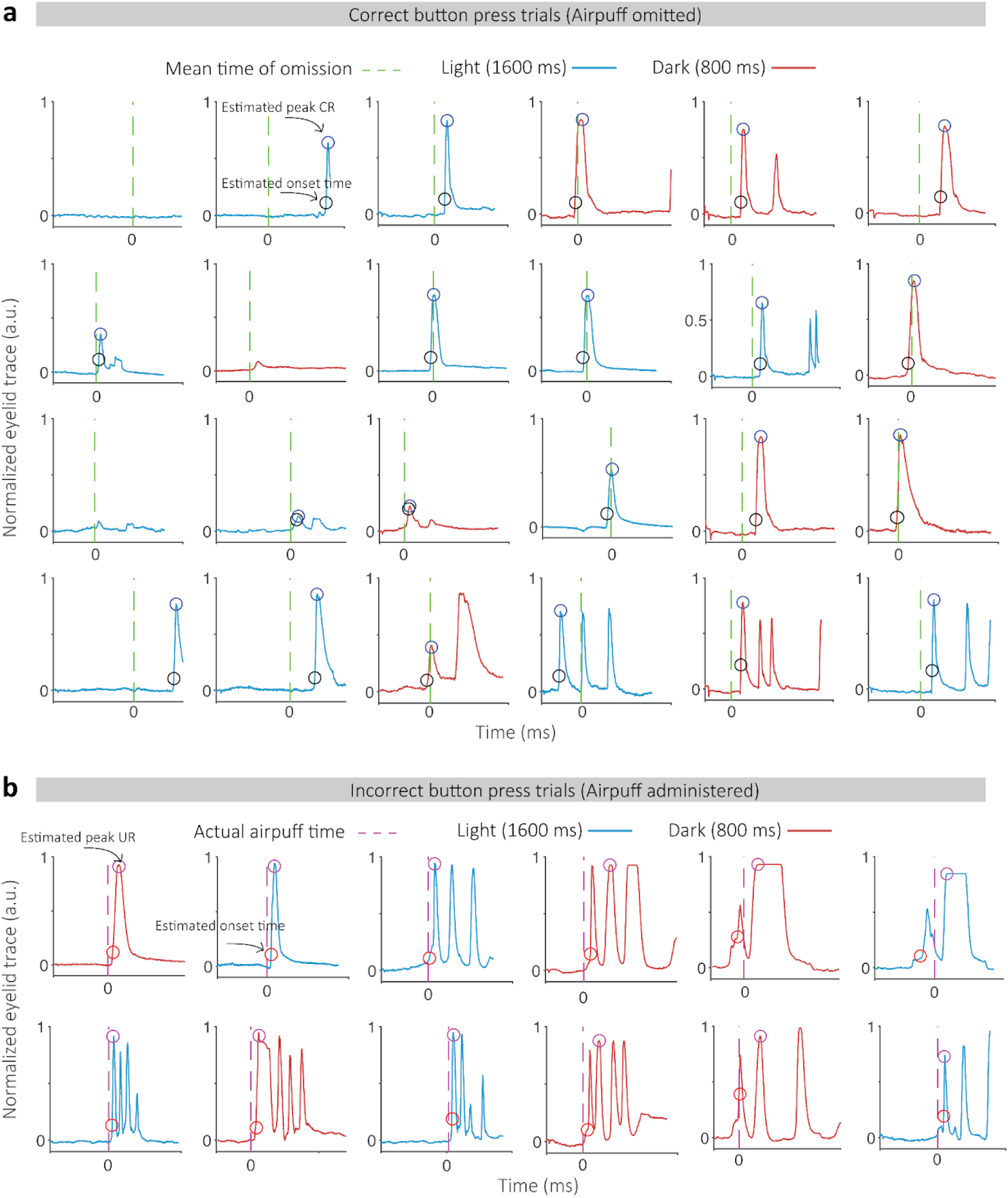
Conditioned and Reflexive eyeblink responses. a) Predictive (conditioned response - CR) eyeblink traces for the Dark (red) and Light (blue) tunnel conditions, aligned to the expected time of the airpuff (green) on correct manual response trials where the airpuff was omitted. The estimates of Peak CR (purple circle) and CR onset (black circles) are indicated. Panels from left to right indicate progression during the course of the experiment from earlier to later stages. b) Reflexive component (unconditioned response UR) of eyeblink responses are shown for the Dark (red) and Light (blue) context conditions aligned to the time of the administered airpuff (purple dashed line) on incorrect manual response trials. UR Onset (red circle) and UR peak (purple circle) are indicated. Panels from left to right indicate progression over the course of the experiment.

**Supplementary figure 2:**
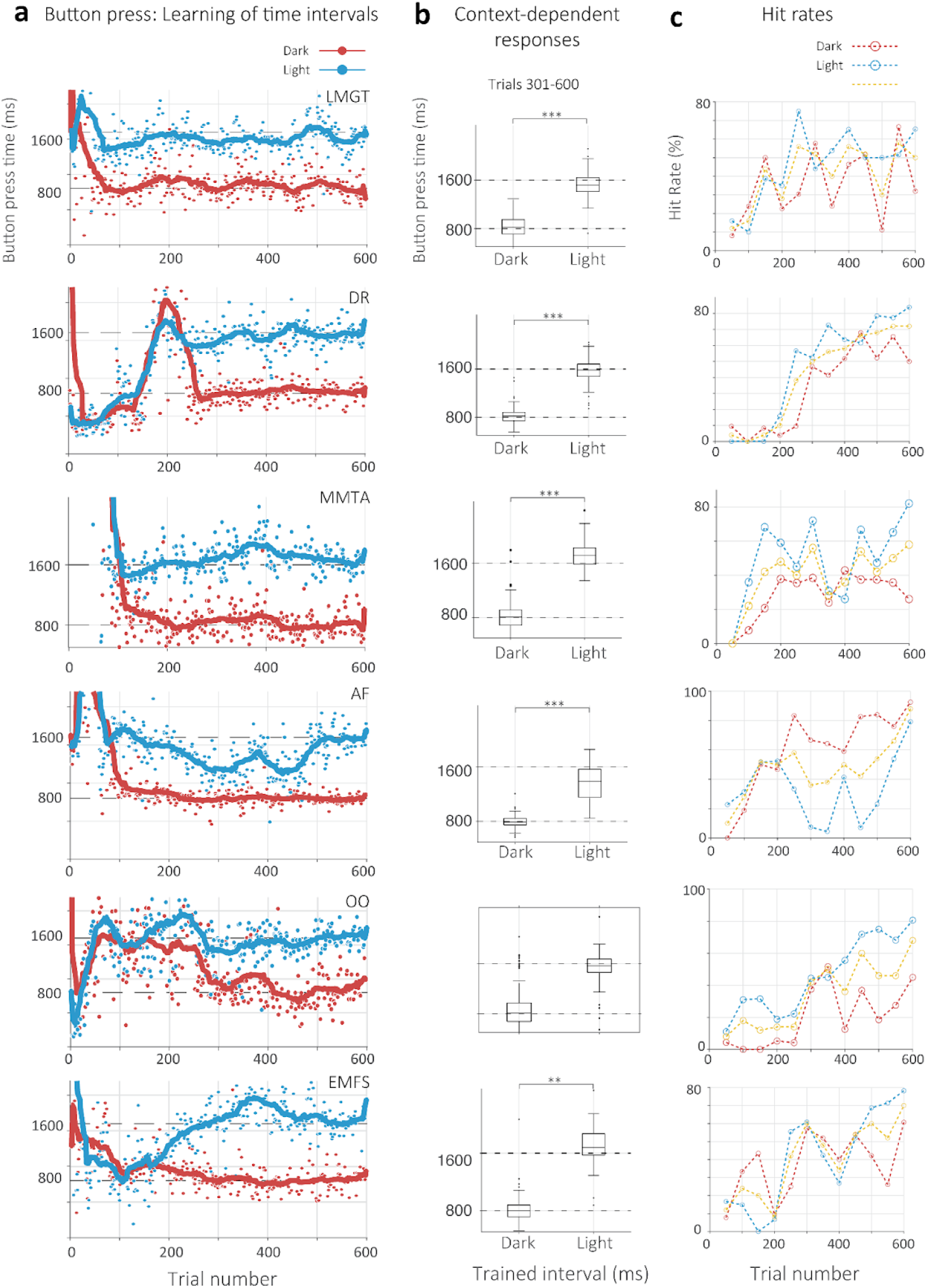
Explicit manual task responses for individuals in naive group: a) Button press time for individual participants for the Dark (red circles) and Light (blue circles) tunnel contexts over the course of the experiments. Solid lines represent moving averages. Black dashed lines indicate the expected time for each context, at 800 and 1600 ms for Dark and Light, respectively. b) Box plots indicating variation in responses for the Dark and Light button press times of each participant evaluated from trial 301-600. Black circles indicate outliers. Error bars indicate quartile ranges. c) Hit rate percentage for each participant over the course of the experiment in bins of 50 trials. Performance is indicated for all trials (yellow dashed line and circles), dark condition (red) and for the light condition (blue).

**Supplementary figure 3:**
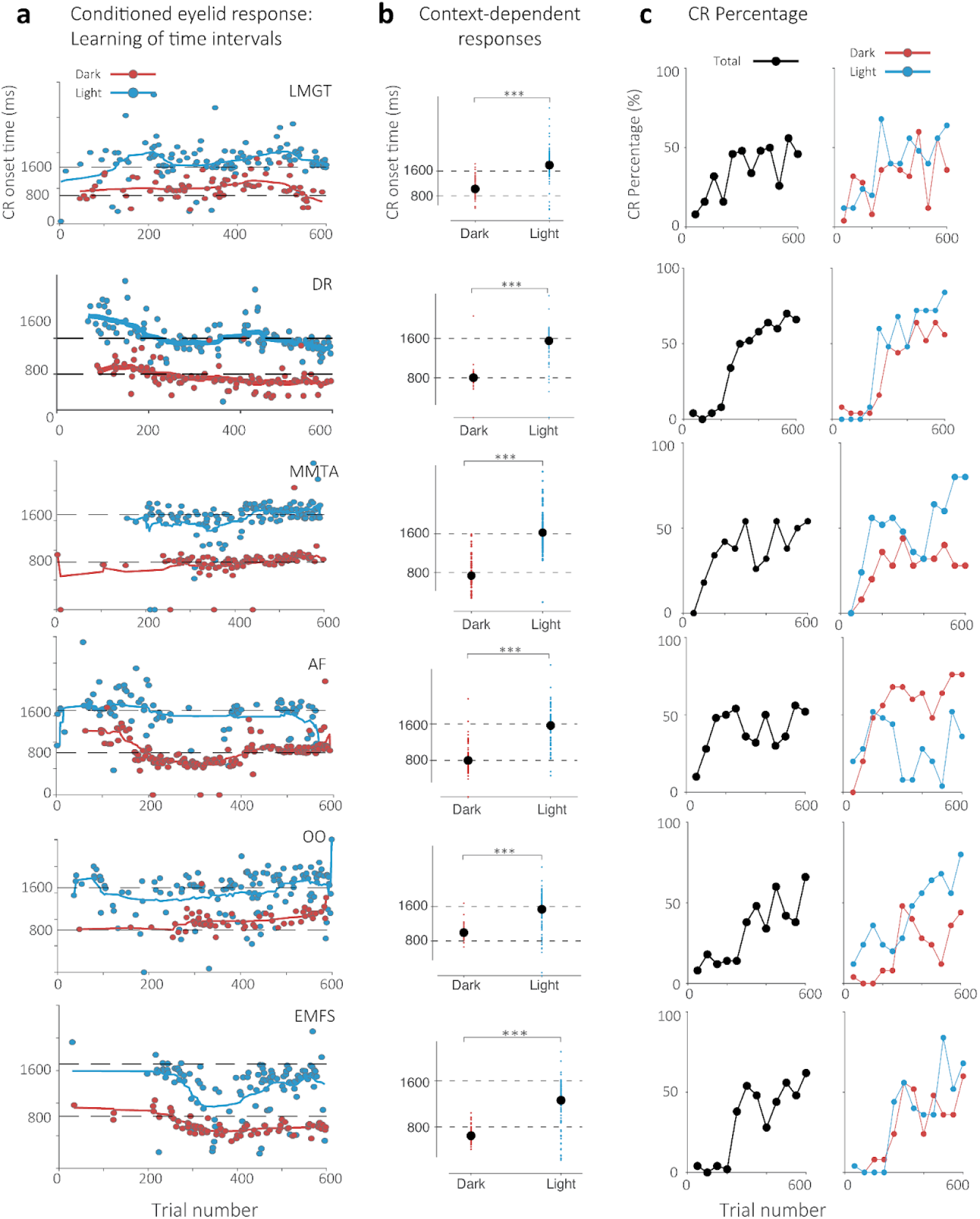
Implicit conditioned eyelid time responses for individuals in naive group: a) Conditioned eyeblink response (CR) onset times evaluated on trials where airpuff was omitted (on correct manual response trials) shown for the Dark (red circles) and Light (blue circles) tunnel contexts. Solid lines represent moving averages. Dashed lines indicate the expected correct manual response for each context for either condition. b) Average CR onset times (black circle) evaluated for all trials. Red and blue circles indicate individual onset times for the Dark and Light condition, respectively. Dashed lines indicate the expected manual response time for Dark (800 ms) and Light (1600 ms) conditions. c) Performance of conditioned responses. CR percentage is shown as a function of number of detected CRs per 50 trials (irrespective of whether they were correct or incorrect manual responses). This measure partly correlates with Hit percentage for manual button press because CRs were only evaluated on correct manual response trials. Left: CR percentage evolution for all trials. Right: CR percentage evolution for the Dark (red) and Light (blue) conditions.

**Supplementary figure 4:**
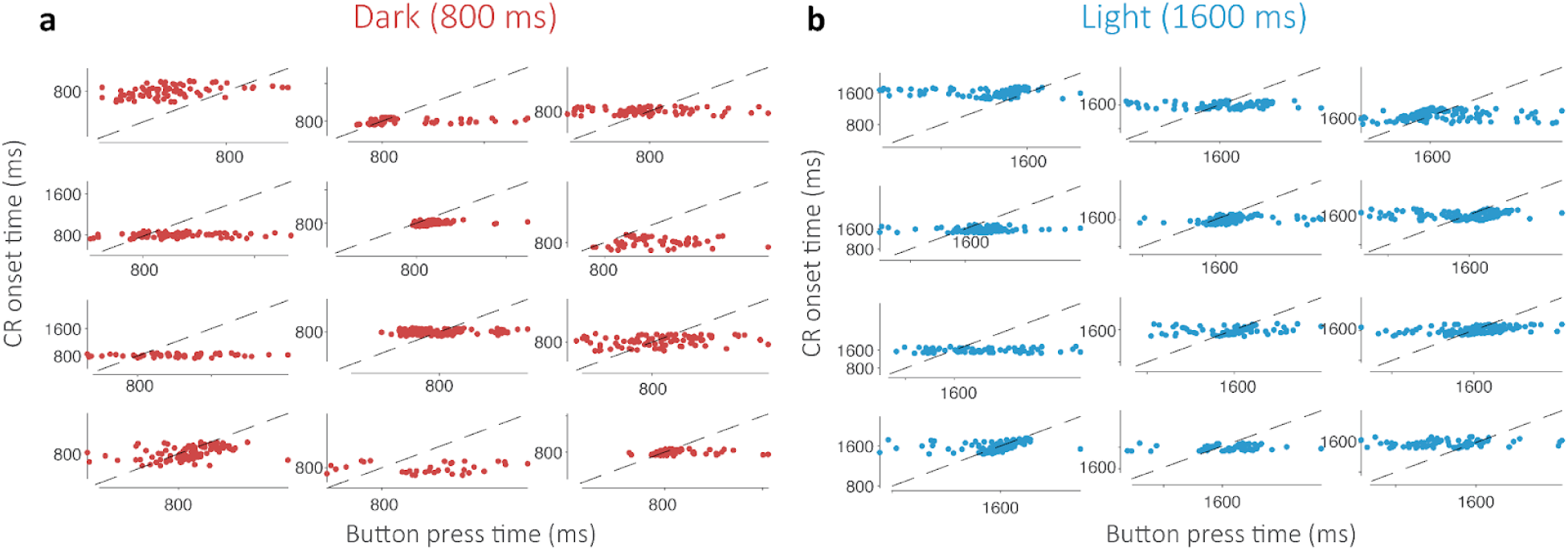
Correlations between CR onset time and button press time. a) Manual button press times are plotted against the CR onset time for individual participants for the Dark (red circles) and Light (blue circles) context condition. Black dashed lines represent unit slope.

**Supplementary figure 5:**
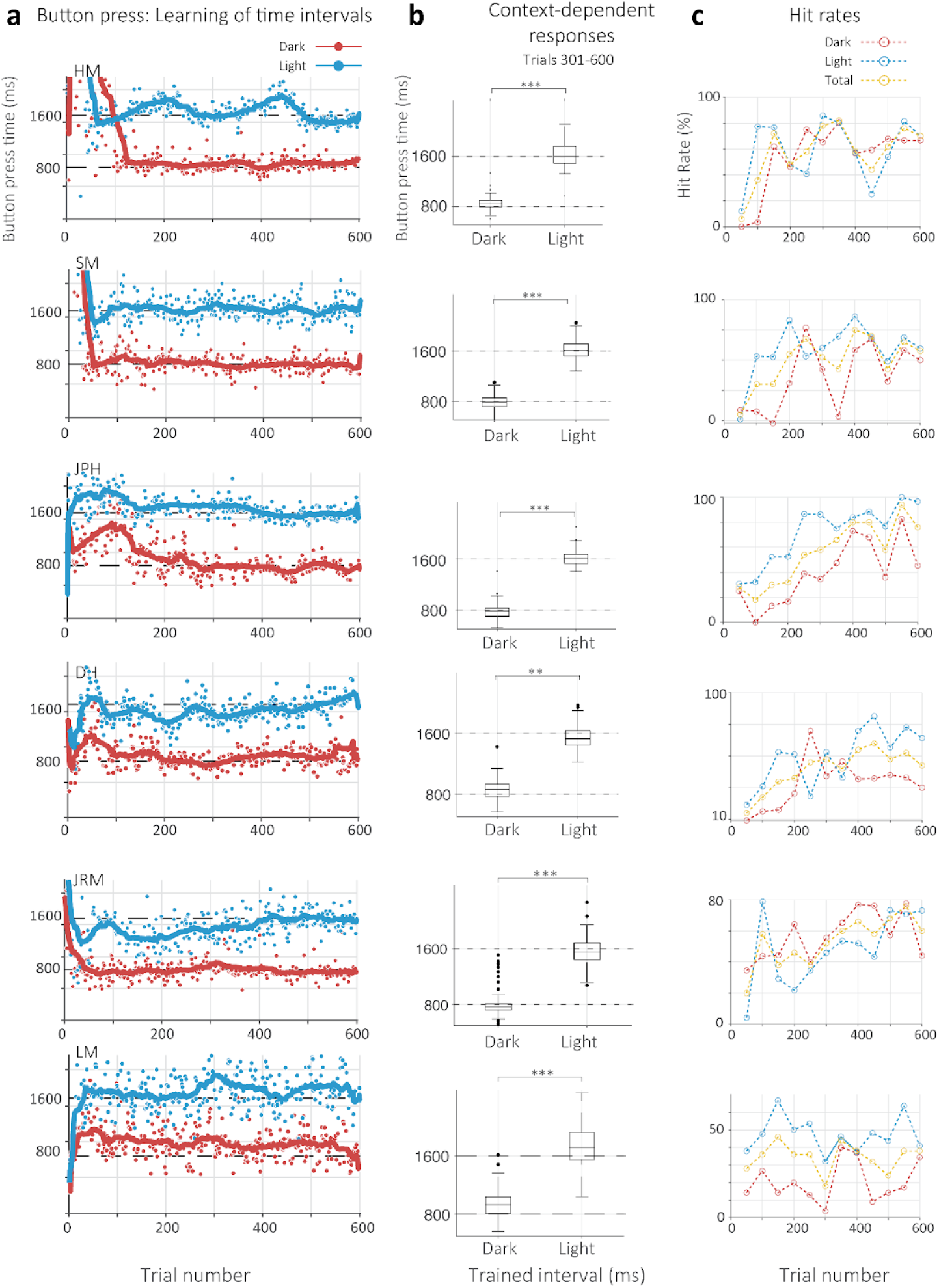
Explicit manual task responses for individuals in strategy group: a) Button press time for individual participants for the Dark (red circles) and Light (blue circles) tunnel contexts over the course of the experiments. Solid lines represent moving averages. Black dashed lines indicate the expected time for each context, at 800 and 1600 ms for Dark and Light, respectively. b) Box plots indicating variation in responses for the Dark and Light button press times of each participant evaluated from trial 301-600. Black circles indicate outliers. Error bars indicate quartile ranges. c) Hit rate percentage for each participant over the course of the experiment in bins of 50 trials. Performance is indicated for all trials (yellow dashed line and circles), dark condition (red) and for the light condition (blue).

**Supplementary figure 6:**
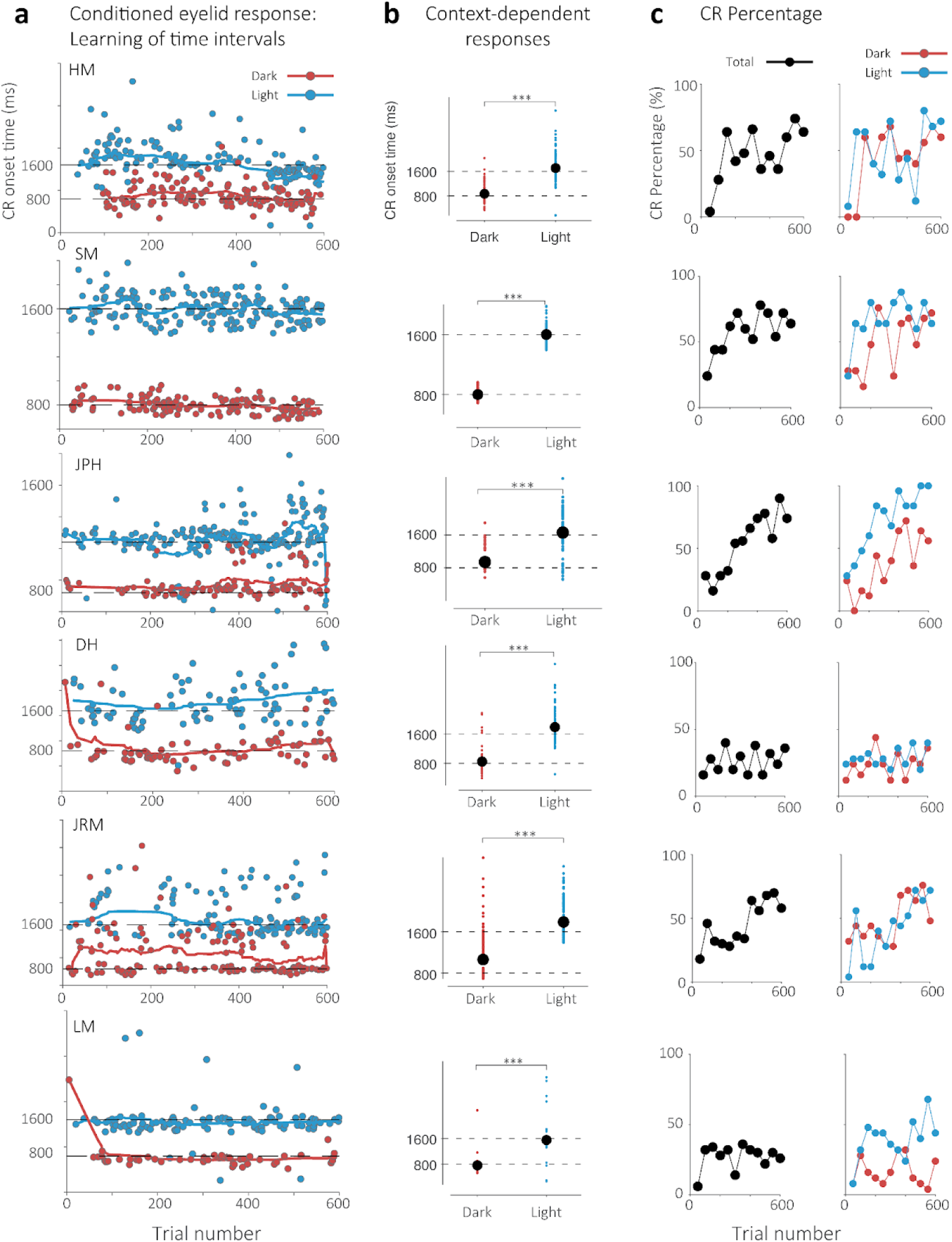
Implicit conditioned eyelid time responses for individuals in strategy group: a) Conditioned eyeblink response (CR) onset times evaluated on trials where airpuff was omitted (on correct manual response trials) shown for the Dark (red circles) and Light (blue circles) tunnel contexts. Solid lines represent moving averages. Dashed lines indicate the expected correct manual response for each context for either condition. b) Average CR onset times (black circle) evaluated for all trials. Red and blue circles indicate individual onset times for the Dark and Light conditions, respectively. Dashed lines indicate the expected manual response time for Dark (800 ms) and Light (1600 ms) conditions. c) Performance of conditioned responses. CR percentage is shown as a function of the number of detected CRs per 50 trials (irrespective of whether they were correct or incorrect manual responses). This measure partly correlates with Hit percentage for manual button press because CRs were only evaluated on correct manual response trials. Left: CR percentage evolution for all trials. Right: CR percentage evolution for the Dark (red) and Light (blue) conditions.

**Supplementary table 1:** ButtonPress statistics for the No Instruction (Naïve) group for participants in Supplementary figure 2. We report statistics of a paired t-test (no tail), with degrees of freedom (df), t statistic (tstat) and p-value for different panels.

| Participant | BoxPlot |  |  | Fraction of correct Trials |  |  |  |  |  |  |  |  |
| --- | --- | --- | --- | --- | --- | --- | --- | --- | --- | --- | --- | --- |
|  |  |  |  | Dark |  |  | Light |  |  | Total |  |  |
|  | Df | tstats | P value | Df | tstats | p | Df | tstats | p | Df | tstats | p |
| LMGT | 298 | -32.6264 | 2.2942e-100 | 11 | 6.5242 | 4.2836e-05 | 11 | 8.3104 | 4.5404e-06 | 11 | 8.8944 | 2.3523e-06 |
| DR | 298 | -33.2178 | 3.4313e-102 | 11 | 4.5931 | 7.7365e-04 | 11 | 4.8907 | 4.7864e-04 | 11 | 4.9556 | 4.3176e-04 |
| MMTA | 298 | -39.8573 | 1.9200e-121 | 11 | 7.3830 | 1.3893e-05 | 11 | 7.3316 | 1.4824e-05 | 11 | 8.2783 | 4.7118e-06 |
| AF | 298 | -24.5370 | 1.6688e-73 | 11 | 7.4784 | 1.2328e-05 | 11 | 5.1732 | 3.0697e-04 | 11 | 8.4437 | 3.8957e-06 |
| OO | 298 | -21.9934 | 2.3025e-64 | 11 | 3.7586 | 0.0032 | 11 | 6.7242 | 3.2682e-05 | 11 | 5.8084 | 1.1784e-04 |
| EMFS | 298 | -35.8361 | 4.7505e-110 | 11 | 7.3605 | 1.4292e-05 | 11 | 5.2420 | 2.7598e-04 | 11 | 6.8274 | 2.8481e-05 |

**Supplementary table 2:** Conditioned Eyeblink response statistics for the No Instruction (Naïve) group for participants in Supplementary figure 3. We report statistics of a paired t-test (no tail), with degrees of freedom (df), t statistic (tstat) and p-value for different panels.

| Participant | CR Time |  |  | CR Percentage |  |  |  |  |  |  |  |  |
| --- | --- | --- | --- | --- | --- | --- | --- | --- | --- | --- | --- | --- |
|  |  |  |  | Dark |  |  | Light |  |  | Total |  |  |
|  | Df | tstats | P value | Df | tstats | p | Df | tstats | p | Df | tstats | p |
| LMGT | 190 | -12.2088 | 1.1054e-25 | 11 | 6.3851 | 5.1860e-05 | 11 | 7.1253 | 1.9292e-05 | 11 | 7.7286 | 9.0551e-06 |
| DR | 233 | -21.0159 | 1.0393e-55 | 11 | 4.7840 | 5.6775e-04 | 11 | 4.6656 | 6.8748e-04 | 11 | 4.9205 | 4.5653e-04 |
| MMTA | 217 | -20.1512 | 1.3299e-51 | 11 | 7.4538 | 1.2712e-05 | 11 | 7.3542 | 1.4407e-05 | 11 | 7.7606 | 8.7101e-06 |
| AF | 239 | -18.6750 | 1.3301e-48 | 11 | 8.1860 | 5.2462e-06 | 11 | 5.7446 | 1.2940e-04 | 11 | 10.0711 | 6.8873e-07 |
| OO | 194 | -9.7262 | 1.8341e-18 | 11 | 4.0829 | 0.0018 | 11 | 6.7963 | 2.9683e-05 | 11 | 5.8004 | 1.1922e-04 |
| EMFS | 188 | -14.1271 | 2.3527e-31 | 11 | 4.7866 | 5.6542e-04 | 11 | 4.2472 | 0.0014 | 11 | 4.7336 | 6.1574e-04 |

**Supplementary table 3:** Accuracy and precision rations for the light and dark conditions for the Naive group.

| Participant | AccuracyRatio Dark | Accuracy Ratio Light | Precision Ratio Dark | Precision Ratio Light |
| --- | --- | --- | --- | --- |
| FSuppl |  |  |  |  |
| 1 | 13,8077 | 1,6027 | 0,1161 | 0,1745 |
| 2 | 1,7091 | 10,25 | 0,0797 | 0,1649 |
| 3 | 0,1908 | 0,6701 | 0,2398 | 0,2634 |
| 4 | 0,9402 | 6,5455 | 0,1613 | 0,138 |
| 5 | 7,625 | 1,711 | 0,1001 | 0,1113 |
| 6 | 0,8571 | 0,9188 | 0,1191 | 0,1305 |
| 7 | 0,9088 | 0,4722 | 0,0846 | 0,196 |
| 8 | 0,8247 | 1,2485 | 0,1414 | 0,2084 |
| 9 | 0,0739 | 0,5982 | 0,0958 | 0,1936 |
| 10 | 0,4588 | 0,2237 | 0,0863 | 0,8293 |
| 11 | 2,4211 | 2,495 | 0,0611 | 0,175 |
| 12 | 0,383 | 0,536 | 0,1363 | 0,1834 |
| 13 | 3,1166 | 3,6262 | 0,1721 | 0,249 |
| 14 | 1,1034 | 0,4835 | 0,2782 | 0,1945 |
| 15 | 0,6684 | 0,4132 | 0,1244 | 0,2332 |
| 16 | 0,0943 | 0,1061 | 0,2681 | 0,2179 |
| 17 | 9,0116 | 4,0979 | 0,1546 | 0,1831 |
| 18 | 1,1325 | 0,9174 | 0,1639 | 0,2333 |
| 19 | 2,5465 | 2,2278 | 0,2442 | 0,2069 |
| 20 | 2,8213 | 4,4335 | 0,0824 | 0,1309 |
| 21 | 3,6706 | 1,5776 | 0,1988 | 0,1895 |
| 22 | 1,5734 | 0,6395 | 0,1454 | 0,1668 |
| 23 | 0,5596 | 0,4451 | 0,1984 | 0,2887 |
| 24 | 1,1561 | 0,898 | 0,0775 | 0,1021 |
| 25 | 0,0211 | 0,2793 | 0,1187 | 0,306 |
| 26 | 4,1628 | 4,8426 | 0,1452 | 0,1546 |
| 27 | 0,8735 | 0,0872 | 0,4571 | 0,201 |
| 28 | 1,8797 | 6,5333 | 0,0841 | 0,1713 |
| 29 | 3,3586 | 12,4028 | 0,1157 | 0,2374 |
| 30 | 4,656 | 2,25 | 0,1175 | 0,1527 |

**Supplementary table 4:**
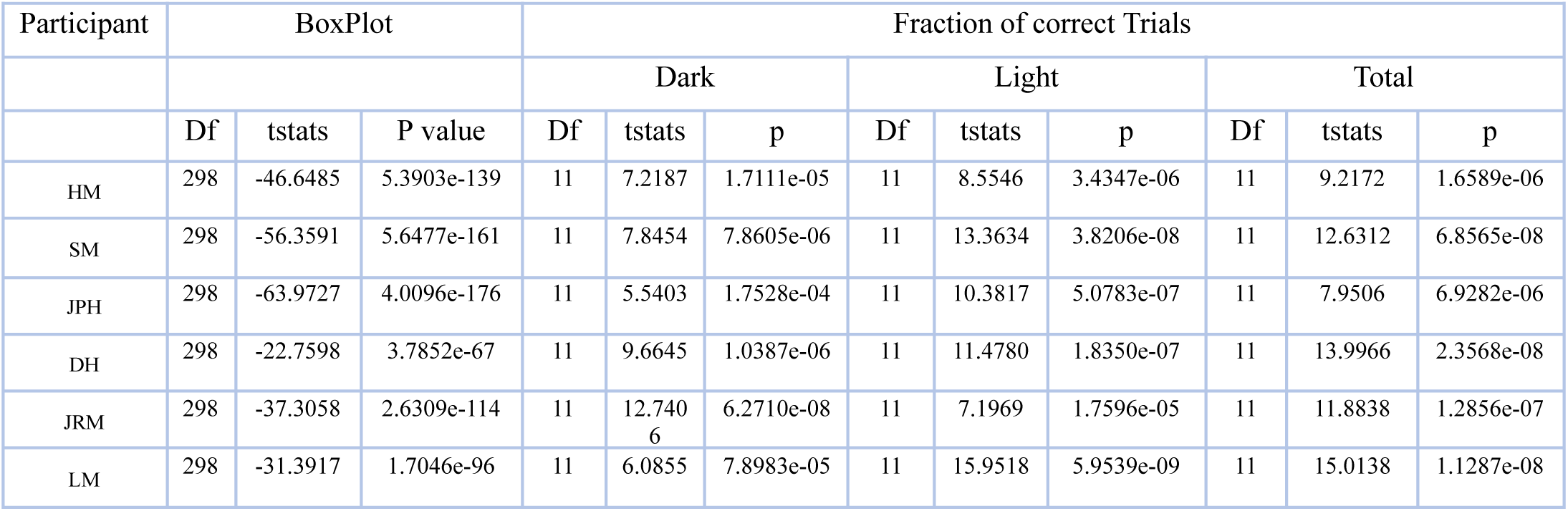
ButtonPress statistics for the Overt Instruction (Strategy) group for participants in Supplementary figure 5. We report statistics of a paired t-test (no tail), with degrees of freedom (df), t statistic (tstat) and p-value for different panels.

**Supplementary table 5:**
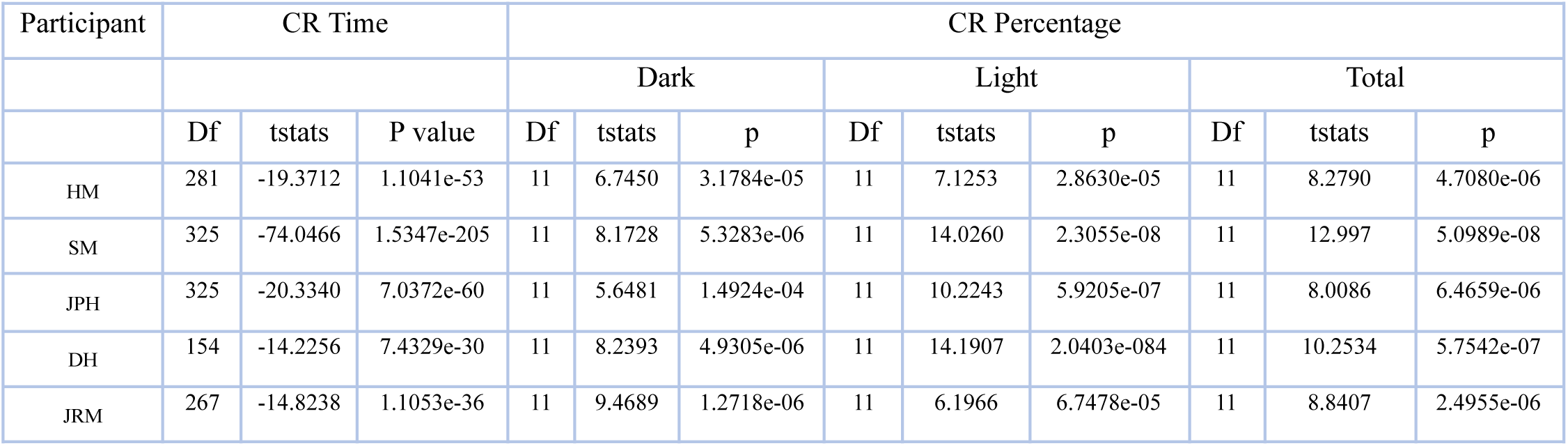
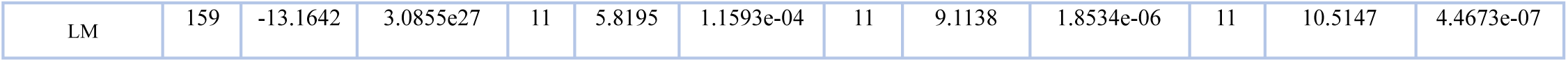
Conditioned eyeblink response statistics for the Overt Instruction (Stategy) group for various participants in Supplementary figure 6. We report statistics of a paired t-test (no tail), with degrees of freedom (df), t statistic (tstat) and p-value for different panels.

| Participant | CR Time |  |  | CR Percentage |  |  |  |  |  |  |  |  |
| --- | --- | --- | --- | --- | --- | --- | --- | --- | --- | --- | --- | --- |
|  |  |  |  | Dark |  |  | Light |  |  | Total |  |  |
|  | Df | tstats | P value | Df | tstats | p | Df | tstats | p | Df | tstats | p |
| HM | 281 | -19.3712 | 1.1041e-53 | 11 | 6.7450 | 3.1784e-05 | 11 | 7.1253 | 2.8630e-05 | 11 | 8.2790 | 4.7080e-06 |
| SM | 325 | -74.0466 | 1.5347e-205 | 11 | 8.1728 | 5.3283e-06 | 11 | 14.0260 | 2.3055e-08 | 11 | 12.997 | 5.0989e-08 |
| JPH | 325 | -20.3340 | 7.0372e-60 | 11 | 5.6481 | 1.4924e-04 | 11 | 10.2243 | 5.9205e-07 | 11 | 8.0086 | 6.4659e-06 |
| DH | 154 | -14.2256 | 7.4329e-30 | 11 | 8.2393 | 4.9305e-06 | 11 | 14.1907 | 2.0403e-084 | 11 | 10.2534 | 5.7542e-07 |
| JRM | 267 | -14.8238 | 1.1053e-36 | 11 | 9.4689 | 1.2718e-06 | 11 | 6.1966 | 6.7478e-05 | 11 | 8.8407 | 2.4955e-06 |
| LM | 159 | -13.1642 | 3.0855e27 | 11 | 5.8195 | 1.1593e-04 | 11 | 9.1138 | 1.8534e-06 | 11 | 10.5147 | 4.4673e-07 |

**Supplementary table 6:**
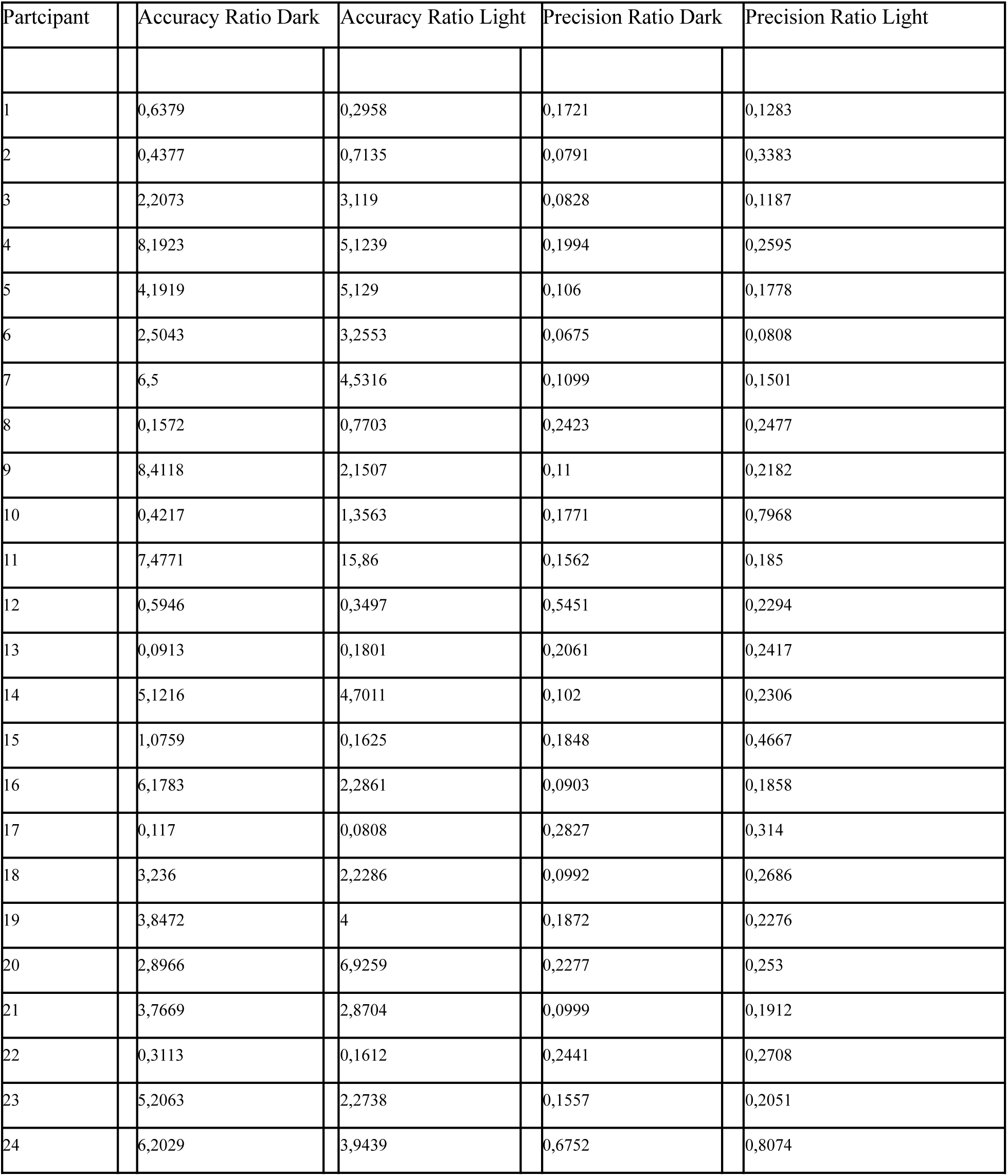
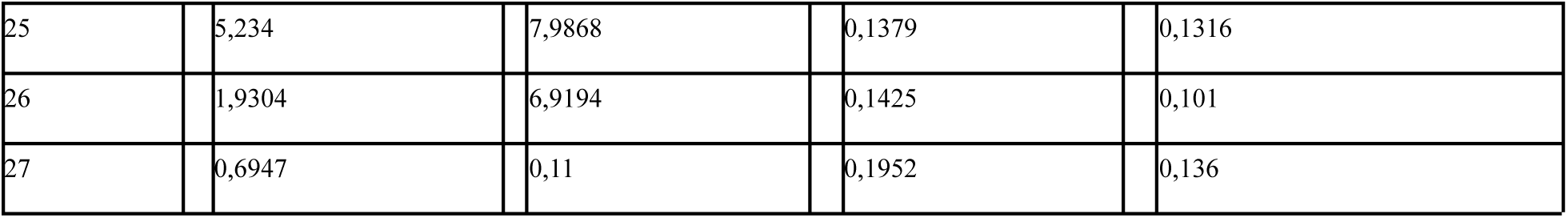
Accuracy and precision rations for the light and dark conditions for the Strategy group.

| Participant | Accuracy Ratio Dark | Accuracy Ratio Light | Precision Ratio Dark | Precision Ratio Light |
| --- | --- | --- | --- | --- |
| 1 | 0,6379 | 0,2958 | 0,1721 | 0,1283 |
| 2 | 0,4377 | 0,7135 | 0,0791 | 0,3383 |
| 3 | 2,2073 | 3,119 | 0,0828 | 0,1187 |
| 4 | 8,1923 | 5,1239 | 0,1994 | 0,2595 |
| 5 | 4,1919 | 5,129 | 0,106 | 0,1778 |
| 6 | 2,5043 | 3,2553 | 0,0675 | 0,0808 |
| 7 | 6,5 | 4,5316 | 0,1099 | 0,1501 |
| 8 | 0,1572 | 0,7703 | 0,2423 | 0,2477 |
| 9 | 8,4118 | 2,1507 | 0,11 | 0,2182 |
| 10 | 0,4217 | 1,3563 | 0,1771 | 0,7968 |
| 11 | 7,4771 | 15,86 | 0,1562 | 0,185 |
| 12 | 0,5946 | 0,3497 | 0,5451 | 0,2294 |
| 13 | 0,0913 | 0,1801 | 0,2061 | 0,2417 |
| 14 | 5,1216 | 4,7011 | 0,102 | 0,2306 |
| 15 | 1,0759 | 0,1625 | 0,1848 | 0,4667 |
| 16 | 6,1783 | 2,2861 | 0,0903 | 0,1858 |
| 17 | 0,117 | 0,0808 | 0,2827 | 0,314 |
| 18 | 3,236 | 2,2286 | 0,0992 | 0,2686 |
| 19 | 3,8472 | 4 | 0,1872 | 0,2276 |
| 20 | 2,8966 | 6,9259 | 0,2277 | 0,253 |
| 21 | 3,7669 | 2,8704 | 0,0999 | 0,1912 |
| 22 | 0,3113 | 0,1612 | 0,2441 | 0,2708 |
| 23 | 5,2063 | 2,2738 | 0,1557 | 0,2051 |
| 24 | 6,2029 | 3,9439 | 0,6752 | 0,8074 |

|  |  |  |  |  |  |  |  |  |
| --- | --- | --- | --- | --- | --- | --- | --- | --- |
| 25 |  | 5,234 |  | 7,9868 |  | 0,1379 |  | 0,1316 |
| 26 |  | 1,9304 |  | 6,9194 |  | 0,1425 |  | 0,101 |
| 27 |  | 0,6947 |  | 0,11 |  | 0,1952 |  | 0,136 |

